## Supporting Information for "Structural basis for higher-order DNA binding by a bacterial transcriptional regulator"

#### SUPPORTING INFORMATION FIGURES

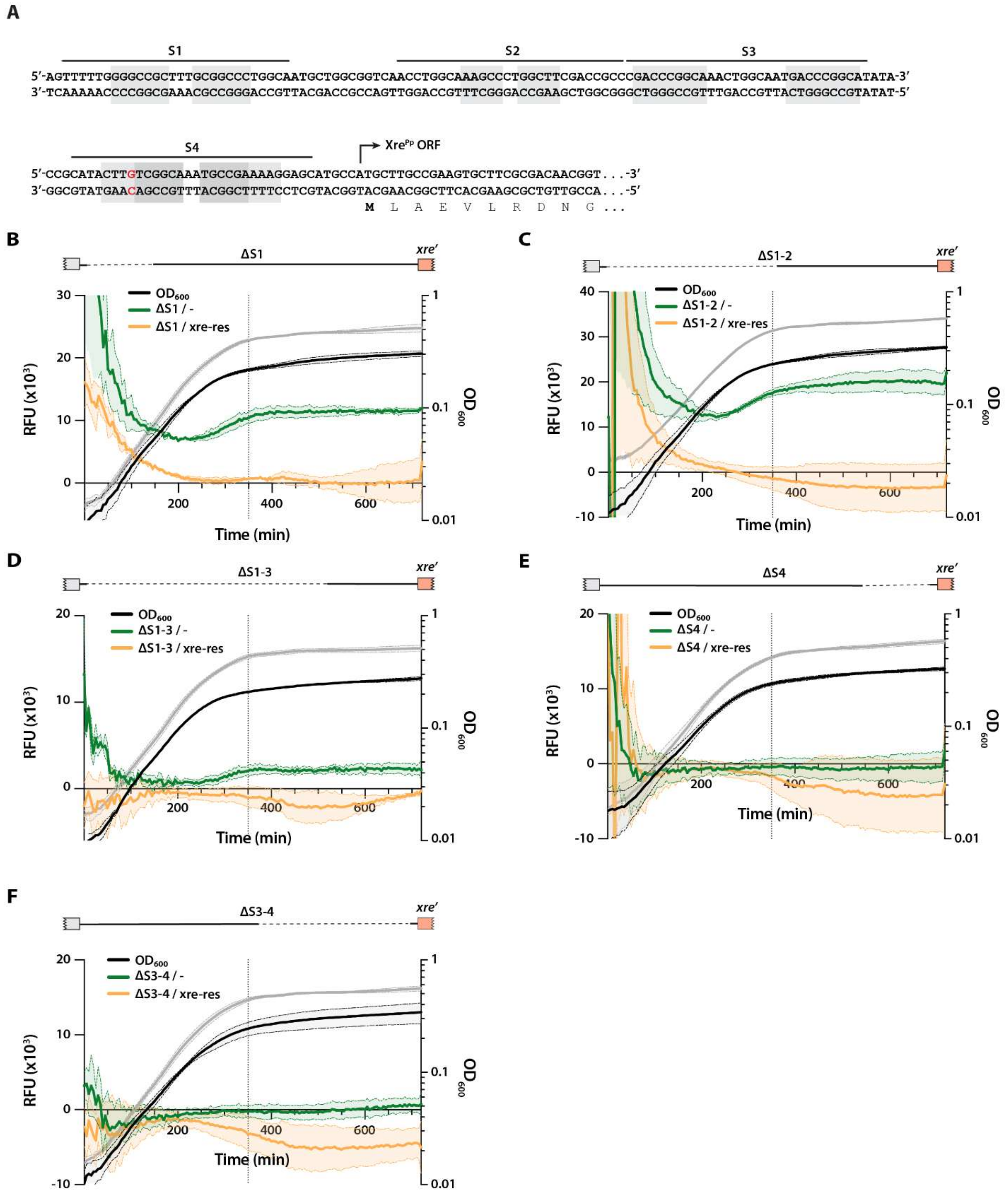

**Fig S1. Determination of the minimal promoter requirements of $P_{XR}$.**

(A) The native DNA sequence of the $P_{XR}$ promoter. The S1, S2, S3, and S4 elements are indicated with lines and light grey boxes marks the repeat. The perfect part of the repeat in S4 is shown with a dark grey box and the mismatch in the first repeat is in red. The first 30-bp of the Xre$^{Pp}$ open reading frame (ORF) is included and marked by the bend arrow, with the N-terminal residues noted below and the start codon in bold. (B)-(F) Promoter activity assays in *E. coli* MG1655. Above, schematic representation of the promoter construct is included above each graph with DNA deletions indicated by dotted lines. Below, promoter activity measured as GFP signal in relative fluorescence units (RFU) during growth without arabinose (green curves) or with arabinose (orange curves) using the promoter reporter plasmid pGH254Kgfp with indicated promoter variations together with empty pBAD33 (-) or pBAD33::*xre-res*$^{Pp}$ (*xre-res*). $OD_{600}$ measurements for cultures without arabinose (grey curves) or with arabinose (black curves) are included. All curves represent mean-of-mean values (line, n = 3) with the standard error of the mean (SEM, shadow). The dotted vertical line indicates the 350 min time point used in **Fig 2B**.

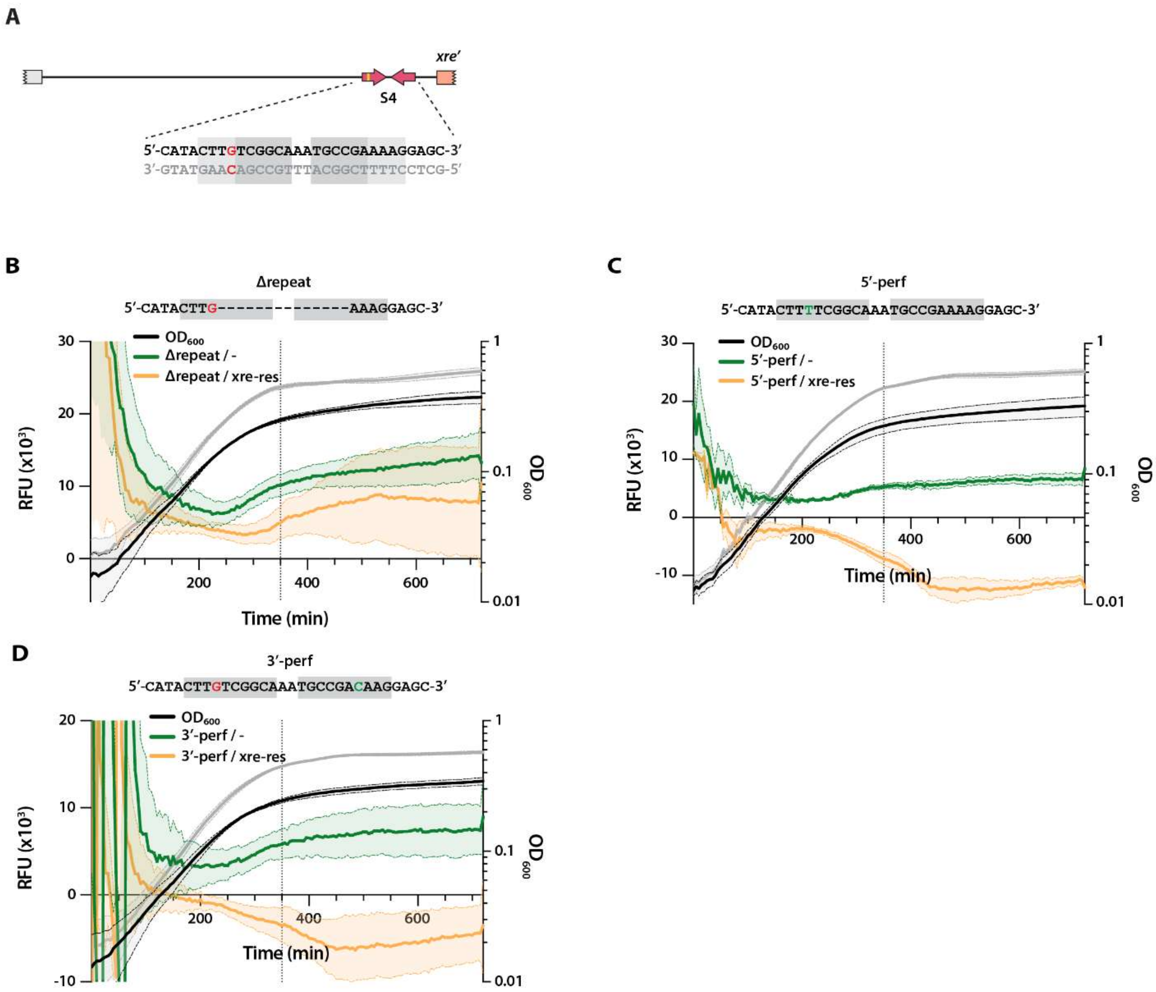


**Fig S2. Functional analysis of the S4 repeat sequence**.

(A) Schematic representation of the $P_{XR}$ promoter. Above, the imperfect inverted repeat (red arrows with mismatch in yellow) in the S4 element is shown. Below, the sequence of the S4 highlighting the inverted repeat (light grey), the mismatch (red), and the perfect part of the repeat (dark grey). (B)-(D) Promoter activity assays in *E. coli* MG1655. Above, schematic representation of the promoter construct with DNA deletions (dotted lines) or modifications (green) indicated. Below, promoter activity measured as GFP signal in RFU during growth without arabinose (green curves) or with arabinose (orange curves) using the promoter reporter pGH254Kgfp with indicated promoter variations together with empty pBAD33 (-) or pBAD33::*xre-res*$^{Pp}$ (*xre-res*). $OD_{600}$ measurements for cultures without (grey curves) or with arabinose (black curves) are included. All curves represent mean-of-mean values (line, n = 3) with the standard error of the mean (SEM, shadow). The dotted vertical line indicates the 350 min time point used in **Fig 2D**.

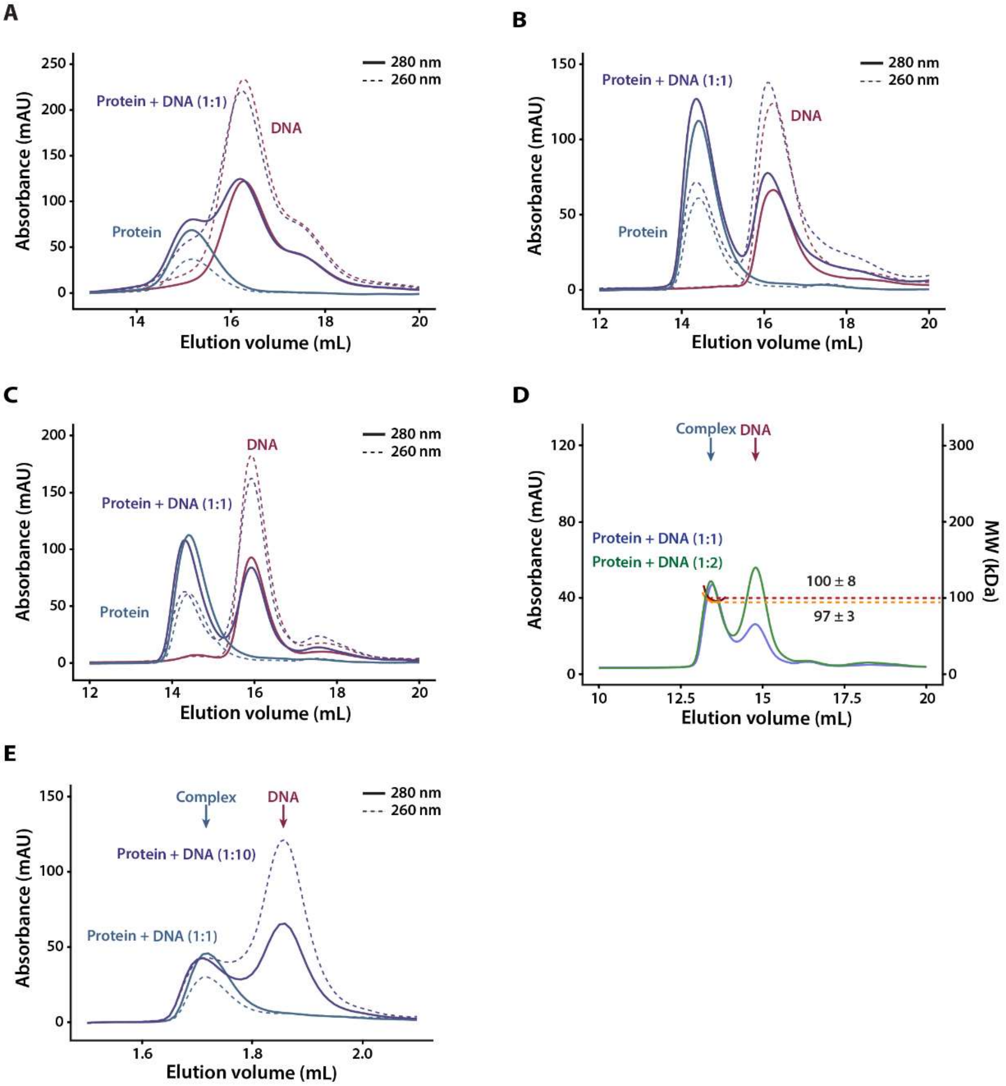


**Fig S3. Xre-RES$^{Pp}$ interacts specifically with S4 in a 1:1 protein complex to DNA duplex ratio**. (A)-(C) Analytical SEC (a-SEC) analysis of Xre-RES$^{Pp}{}_{His6}$ binding to S1 dsDNA (A), S2 dsDNA (B), or S3 dsDNA (C). Chromatograms show elution profiles for Xre-RES$^{Pp}{}_{His6}$ (Protein, green), DNA (red), or DNA mixed with protein in a 1:1 ratio (Protein + DNA 1:1, purple). The full line shows 280 nm absorbance, while the dotted line is 260 nm absorbance. (D) SEC-MALS analysis of the change in mass of Xre-RES$^{Pp}{}_{His6}$ mixed with S4 dsDNA. The chromatogram shows the elution profiles for the 1:1 (blue) or the 1:2 (green) Xre-RES$^{Pp}$ complex to S4 DNA duplex ratios, as well as the masses of the peaks for the 1:1 (yellow dotted line) and 1:2 (red dotted line) ratios. The average masses corresponding to the middle of the peaks were determined to 97 ± 3 (1:1 ratio) and 100 ± 8 kDa (1:2

ratio). (E) High resolution a-SEC analysis of Xre-RES$^{Pp}{}_{His6}$ mixed with S4 dsDNA in a 1:1 (green) or 1:10 (purple) ratio. Full lines show 280 nm and dotted lines 260 nm absorbance.

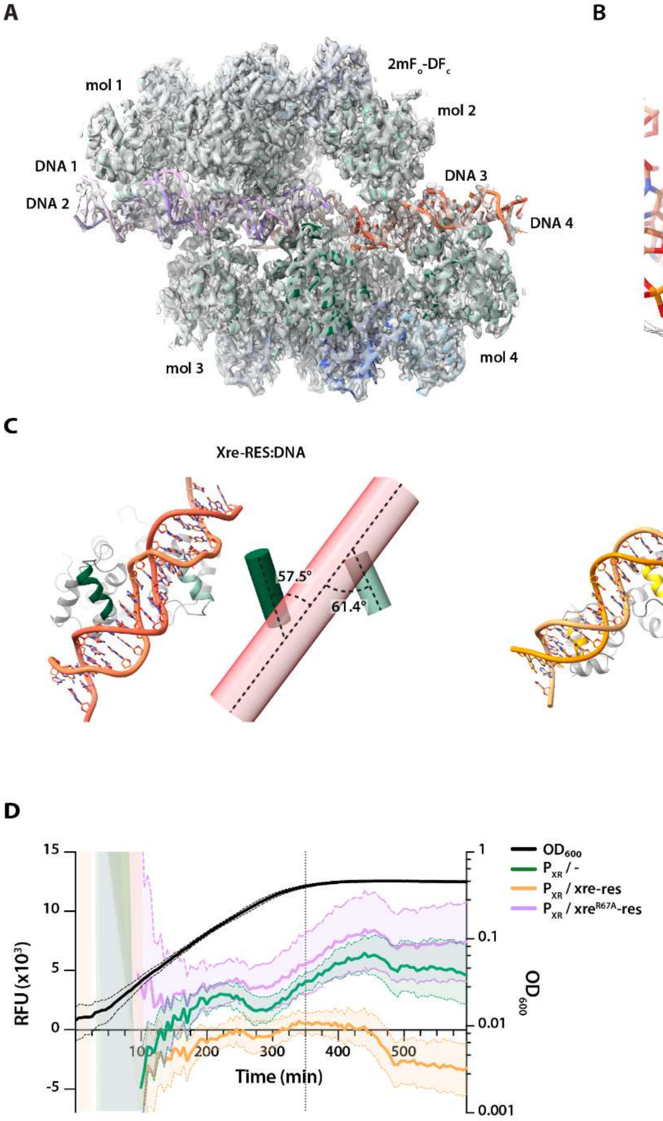


**Fig S4. Crystal structure of the DNA-bound Xre-RES$^{Pp}$ complex.**

(A) The contents of the asymmetric unit (ASU) with electron density (2mFo-Fc, grey) contoured at 1 σ. The ASU contains four copies of the Xre-RES hexamer (mol 1-mol 4) and four copies of S4 DNA (DNA 1-DNA 4). The molecules (protein and DNA) are related through two-fold symmetry with protein in blue/green (lighter for symmetry-related molecules) and DNA in red (for DNA 1 and 3) and purple (for DNA 2 and 4). (B) A close-up view of Xre-RES molecule 1 (chains A-F) and DNA 1 (chain a) showing the high-resolution features of the map, with interacting sidechains Arg60, Arg67, Thr68, and Arg72 in sticks. (C) A focused view of the HtH domains interacting with DNA for Xre-RES (green, left) and the 434 Cro repressor from the lambda phage (PDB 1RPE) (1) (yellow, right).

In each case, the structure is shown in cartoon with the HTH recognition helices colored (left) next to schematic figures showing centroid cylinder as defined by the helices (right) and including the angle between the recognition helices (green, yellow) and the DNA (faint red) (D) Promoter activity assays in *E. coli* MG1655. Activity is measured as GFP signal in RFU during growth for cultures with arabinose, using the pGH254Kgfp::$P_{XR}$ reporter plasmid combined with either empty pBAD33 (green curve), pBAD33::*xre-res*$^{Pp}$ (orange curve) or pBAD33::*xre*$^{R67A}$-*res*$^{Pp}$ (purple curve). $OD_{600}$ measurements for the pGH254Kgfp::$P_{XR}$/pBAD33 culture is included (black curve). Curves show mean-of-mean values (line, n = 3) with SEM (shadow). The dotted vertical line indicates the 350 min time point.

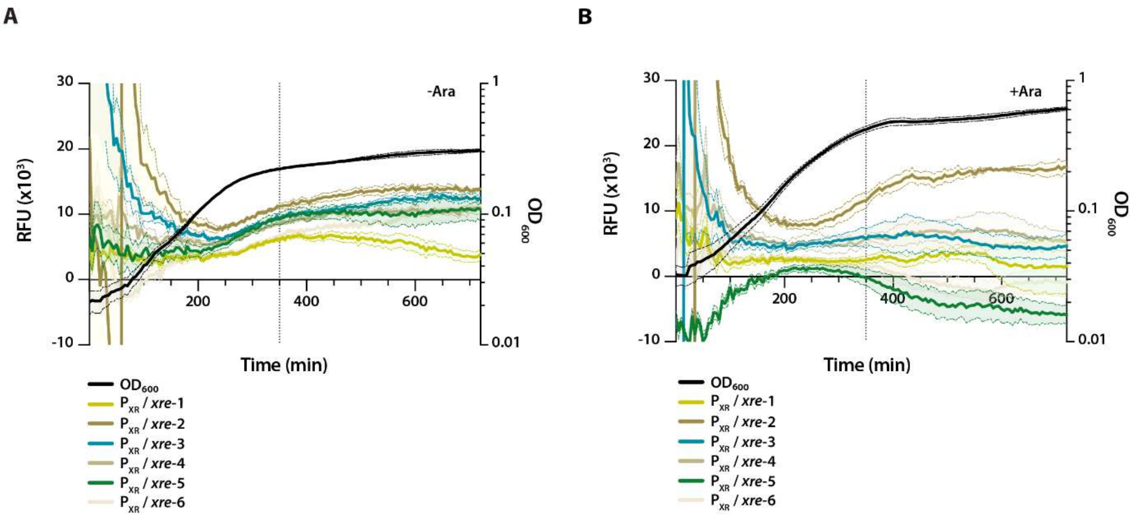


**Fig S5. Expression of *xre*$^{\text{Pp}}$ alone only partially represses transcription from $P_{XR}$.**

(A) and (B) Promoter activity assays in *E. coli* MG1655. Activity is measured as GFP signal in RFU during growth for cultures without arabinose (A) or with arabinose (B), using the pGH254Kgfp::$P_{XR}$ reporter plasmid ($P_{XR}$) combined with pBAD33::*xre* (*xre*). The colors represent six different biological replicates (n = 6), highlighting the variability in GFP expression upon *xre*$^{\text{Pp}}$ expression (B). $OD_{600}$ measurements of one of the biological replicates are included (black curve). All curves represent mean-of-mean values (line, n = 6) with SEM (shadow). The dotted vertical line indicates the 350 min time point used in **Fig 5A**.

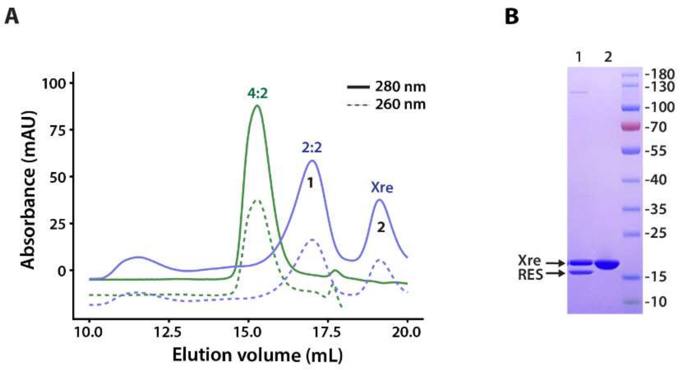


**Fig S6. Isolation of the 2:2 Xre-RES$^{Pp}$ complex**.

(A) Analytical SEC (a-SEC) analysis. Chromatograms show elution profiles for purified the Xre-RES$^{Pp}{}_{His6}$ complex (4:2, green) and the Xre$_{NHis6}$-RES$^{Pp}$ complex (2:2 and free Xre antitoxin, blue). Full line shows 280 nm absorbance, while dotted line is 260 nm absorbance. The identity of peak 1 and 2 was confirmed by SDS-PAGE analysis (B).

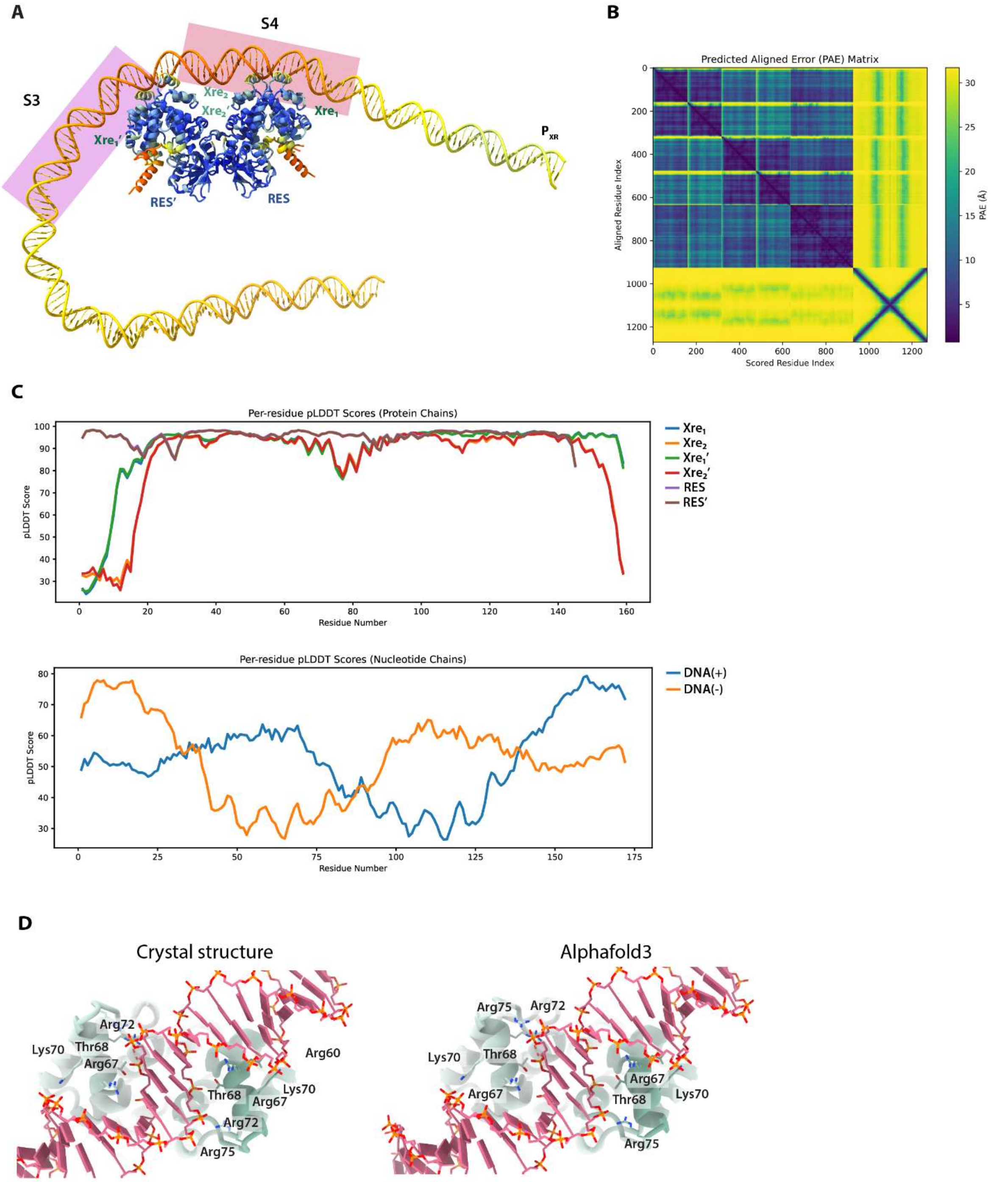


**Fig S7. Alphafold 3 prediction of the interaction between Xre-RES and the full promoter.**

(A) The AlphaFold3 predicted structure of the hexameric $Xre_4RES_2$ complex bound to a 172 bp double stranded DNA segment covering to the promoter and intergenic region between *xre* and the upstream gene in the *Pseudomonas putida* KT2440 genome ($P_{XR}$), the prediction is colored by pLDDT score and regions corresponding to S3 and S4 are highlighted. (B) Predicted Aligned Error (PAE) plot corresponding to the AlphaFold3 prediction. (C) A plot of pLDDT values as a function of residue number for the AlphaFold3 prediction of the protein (top) and the DNA segment (bottom). (D) A comparison between the DNA recognition site from the crystal structure (left) and the AlphaFold3 prediction (right), residues withing 3.8 Å of the DNA helix are shown in sticks and labeled.

### SUPPORTING INFORMATION TABLES

#### S1 Table. Promoter DNA sequences

| | |
|---|---|
| $P_{XR}$ | AGTTTTTGGGGCCGCTTTGCGGCCCTGGCAATGCTGGCGGTCAACCTGGCAAAGCCCTGGCTTCGACCGCCCGACCCGGCAAACTGGCAATGACCCGGCATATACCGCATACTTGTCGGCAAATGCCGAAAAGGAGCATGCC**ATG**CTTGCCGAAGTGCTTCGCGACAACGGT |
| $P_{XR}\Delta$S1 | TGGCGGTCAACCTGGCAAAGCCCTGGCTTCGACCGCCCGACCCGGCAAACTGGCAATGACCCGGCATATACCGCATACTTGTCGGCAAATGCCGAAAAGGAGCATGCC**ATG**CTTGCCGAAGTGCTTCGCGACAACGGT |
| $P_{XR}\Delta$S1-2 | CGACCCGGCAAACTGGCAATGACCCGGCATATACCGCATACTTGTCGGCAAATGCCGAAAAGGAGCATGCC**ATG**CTTGCCGAAGTGCTTCGCGACAACGGT |
| $P_{XR}\Delta$S1-3 | ATACCGCATACTTGTCGGCAAATGCCGAAAAGGAGCATGCC**ATG**CTTGCCGAAGTGCTTCGCGACAACGGT |
| $P_{XR}\Delta$S4 | AGTTTTTGGGGCCGCTTTGCGGCCCTGGCAATGCTGGCGGTCAACCTGGCAAAGCCCTGGCTTCGACCGCCCGACCCGGCAAACTGGCAATGACCCGGCATATACCG---ATGCC**ATG**CTTGCCGAAGTGCTTCGCGACAACGGT |
| $P_{XR}\Delta$3-4 | AGTTTTTGGGGCCGCTTTGCGGCCCTGGCAATGCTGGCGGTCAACCTGGCAAAGCCCTGGCTTCGACCGCC---ATGCC**ATG**CTTGCCGAAGTGCTTCGCGACAACGGT |
| $P_{XR}\Delta$repeat | AGTTTTTGGGGCCGCTTTGCGGCCCTGGCAATGCTGGCGGTCAACCTGGCAAAGCCCTGGCTTCGACCGCCCGACCCGGCAAACTGGCAATGACCCGGCATATACCGCATACTTG---AAAGGAGCATGCC**ATG**CTTGCCGAAGTGCTTCGCGACAACGGT |
| $P_{XR}$-5’perfect | AGTTTTTGGGGCCGCTTTGCGGCCCTGGCAATGCTGGCGGTCAACCTGGCAAAGCCCTGGCTTCGACCGCCCGACCCGGCAAACTGGCAATGACCCGGCATATACCGCATACTTTTCGGCAAATGCCGAAAAGGAGCATGCC**ATG**CTTGCCGAAGTGCTTCGCGACAACGGT |
| $P_{XR}$-3’perfect | AGTTTTTGGGGCCGCTTTGCGGCCCTGGCAATGCTGGCGGTCAACCTGGCAAAGCCCTGGCTTCGACCGCCCGACCCGGCAAACTGGCAATGACCCGGCATATACCGCATACTTGTCGGCAAATGCCGACAAGGAGCATGCC**ATG**CTTGCCGAAGTGCTTCGCGACAACGGT |

The four repeats in the promoter are indicated by colour (S1, yellow; S2, green; S3, purple; S4, brown) with the repeat underlined and the imperfect palindromic part in italics. Dashes (-) indicate deletions and the start codon of the xre gene is shown with a boldface ATG. Substitutions in relation to the wild type sequence ($P_{XR}$, top) are shown with red letters.

**S2 Table. Strains and plasmids used in this study.**

| **Strains** | **Genotype or description** | **Source** |
|---|---|---|
| MG1655 | *E. coli* K-12 *F– λ– ilvG– rfb-50 rph-1* | Laboratory collection |
| BL21(DE3) | *E. coli* B $F^{-}$ *dcm ompT hsdS*(*rB*⁻ *mB*⁻) *gal* | Agilent |
| **Plasmids** | | |
| pUC57::*xre-res*$^{Pp}$ | *P. putida* KT2440 *xre-res* locus including 227bp upstream/60bp downstream | (2) |
| pBAD33 | p15, *araC*, $P_{BAD}$, $Cm^r$ | (3) |
| pBAD33::$xre_{opSD}^{Pp}$ | *P. putida xre* with an optimized Shine-Dalgarno (opSD)[1] | This work |
| pBAD33::$xre_{naSD}$-$res^{Pp}$ | *P. putida xre-res* locus with a native SD (naSD) | This work |
| pBAD33::$xre_{naSD}^{R67A}$-$res^{Pp}$ | *P. putida* $xre^{R67A}$-*res* locus with a native SD (naSD) | This work |
| pGH254K | Mini-R1, *lacZYA* transcriptional fusion vector, $Kan^r$ | (4) |
| pGH254Kgfp | Mini-R1, *gfp* transcriptional fusion vector, $Kan^r$ | This work |
| pGH254Kgfp::$P_{XR}$ | Transcriptional fusion of $P_{XR}$-*xre'* to *gfp* | This work |
| pGH254Kgfp::$P_{XR}$Δ1 | Deletion of Sequence 1 in $P_{XR}$ (bp 1-34) | This work |
| pGH254Kgfp::$P_{XR}$Δ1-2 | Deletion of Sequence 1-2 in $P_{XR}$ (bp 1-71) | This work |
| pGH254Kgfp::$P_{XR}$Δ1-3 | Deletion of Sequence 1-3 in $P_{XR}$ (bp 1-101) | This work |
| pGH254Kgfp::$P_{XR}$Δ4 | Deletion of Sequence 4 in $P_{XR}$ (bp 107-137) | This work |
| pGH254Kgfp::$P_{XR}$Δ3-4 | Deletion of Sequence 3-4 in $P_{XR}$ (bp 71-137) | This work |
| pGH254Kgfp::$P_{XR}$Δrepeat | Deletion of perfect part of repeat in Sequence 4 in $P_{XR}$ (bp 115-129) | This work |
| pGH254Kgfp::$P_{XR}$5'perf | Perfect 5'-repeat in Sequence 4 (substitution G115T) | This work |
| pGH254Kgfp::$P_{XR}$3'perf | Perfect 3'-repeat in Sequence 4 (substitution A130C) | This work |
| pET-29b(+) | *ori*ColE1, $Km^r$, T7 promoter | Twist Bioscience |
| pET-29b(+)::$xre_{CHis6}^{Pp}$ | *P. putida xre* with a C-terminal $His_6$-tag | Twist Bioscience |
| pETDuet-1 | $Amp^r$, *lacI*, T7 promoter | Novagene |
| pETDuet::$res_{NHis6}$-$xre^{Pp}$ | *P. putida res* in MCSI, adding N-terminal $His_6$-tag, and *xre* in MCSII | This work |
| pETDuet::$xre_{NHis6}$-$res^{Pp}$ | *P. putida xre* in MCSI, adding N-terminal $His_6$-tag, and *res* in MCSII | This work |

The names of the strains and plasmids are listed together with a genotype or description and the source. Detailed information about the plasmid constructions can be found in **Appendix Supplementary Methods**. [1]The optimized (op) Shine-Dalgarno (SD) sequence (TAAGGAGGAAATTAA) was included in 5' end of the construct and composed of a strong SD (TAAGGAGG) (5) and additional A's and T's (AAATTAA) to optimize translation.

**S3 Tabel. DNA oligonucleotides used in this study**.

| **Oligonucleotide** | **Sequence (5’-3’)** |
|---|---|
| xre_opSD_Fw | CCCCAGGTACCTAAGGAGGAAATTAAATGCTTGCCGAAGTGCTTCG |
| xre_Rv | AAAAAGTCGACTCACAGGCCATAGCCCTCG |
| xre_naSD_Fw | AAAACGGTACCCAAATGCCGAAAAGGAGCATGCC |
| res_Rv | AAAAAGTCGACCTAGAACAGGCGGCTGTCC |
| xre(R67A)_Fw | CATCCCGCTGGCGACCCTCAAATCCC |
| xre(R67A)_Rv | ATCTGGTCGCGCTCC |
| pGH254K_gib_Fw | GATGAACTATACAAATAAATATTATAAAAATTGCCTGATACGCTGCG |
| pGH254K_gib_Rv | GTTCTTCTCCTTTACTCATAAGCTGTTTCCTGTGTGATAAAGAAAGT |
| GFPmut2_gib_Fw | TTATCACACAGGAAACAGCTTATGAGTAAAGGAGAAGAACTTTTCACTGGAG |
| GFPmut2_gib_Rv | TATCAGGCAATTTTTATAATATTTATTTGTATAGTTCATCCATGCCATGTG |
| pGH254Kgfp_T6613_Fw | AGGAAACAGCTATGAGTAAAGGAG |
| pGH254Kgfp_T6613_Rv | GTGTGATAAAGAAAGTTAAAATG |
| PXR_Fw | AACCCGAATTCAGTTTTTGGGGCCGCTTTGC |
| PXR_Rv | AAAAAGGATCCACCGTTGTCGCGAAGCACTTC |
| PXRΔS1_Fw | TGGCGGTCAACCTGGCAA |
| PXRΔS1_Rv | GAATTCAGTTTGTAGAAACGCAAAAAGGC |
| PXRΔS1-2_Fw | CGACCCGGCAAACTGGCA |
| PXRΔS1-2_Rv | GAATTCAGTTTGTAGAAACGCAAAAAGGC |
| PXRΔS1-3_Fw | ATACCGCATACTTGTCGG |
| PXRΔS1-3_Rv | GAATTCAGTTTGTAGAAACGC |
| PXRΔS4_Fw | ATGCCATGCTTGCCGAAG |
| PXRΔS4_Rv | CGGTATATGCCGGGTCATT |
| PXRΔS3-4_Fw | ATGCCATGCTTGCCGAAGTGC |
| PXRΔS3-4_Rv | GGCGGTCGAAGCCAGGGC |
| PXRΔrepeat_Fw | AAAGGAGCATGCCATGCT |
| PXRΔrepeat_Rv | CAAGTATGCGGTATATGCC |
| PXR-5’perf_Fw | CCGCATACTTTTCGGCAAATG |
| PXR-5’perf_Rv | TATATGCCGGGTCATTGC |
| PXR-3’perf_Fw | CAAATGCCGACAAGGAGCATG |
| PXR-3’perf_Rv | CCGACAAGTATGCGGTAT |

| | |
|---|---|
| res_mcsI_Fw | AAACCGGATCCGTGATTTTGTGGCGAATCAGCG |
| res_mcsI_Rv | AAAAAGTCGACCTAGAACAGGCGGCTGTCCG |
| xre_mcsII_Fw | AACCCCATATGATGCTTGCCGAAGTGCTTCG |
| xre_mcsII_Rv | AAAAACTCGAGTCACAGGCCATAGCCCTCG |
| xre_mcsI_Fw | AAAAAGGATCCGCTTGCCGAAGTGCTTCGCGAC |
| xre_mcsI_Rv | AAAAAGTCGACTCACAGGCCATAGCCCTCG |
| res_mcsII_Fw | AGGGGCATATGATTTTGTGGCGAATCAGCG |
| res_mcsII_Rv | AAAAACTCGAGCTAGAACAGGCGGCTGTCC |
| S1_fwd | TTTTTGGGGCCGCTTTGCGGCCCTGGCA |
| S1_rev | TGCCAGGGCCGCAAAGCGGCCCCAAAAA |
| S2_fwd | ACCTGGCAAAGCCCTGGCTTCGACCGCC |
| S2_rev | GGCGGTCGAAGCCAGGGCTTTGCCAGGT |
| S3_fwd | CGACCCGGCAAACTGGCAATGACCCGGCAT |
| S3_rev | ATGCCGGGTCATTGCCAGTTTGCCGGGTCG |
| S4_fwd | CATACTTGTCGGCAAATGCCGAAAAGGAGC |
| S4_rev | GCTCCTTTTCGGGATTTGCCGACAAGTATG |
| 5'perf_fwd | CATACTTTTCGGCAAATGCCGAAAAGGAGC |
| S4_5'perf_rev | GCTCCTTTTCGGGATTTGCCGAAAAGTATG |
| S4_3'perf_fwd | CATACTTGTCGGCAAATGCCGACAAGGAGC |
| S4_3'perf_rev | GCTCCTTGTCGGGATTTGCCGACAAGTATG |
| S4_5'flip_fwd | CATAGGCTGTTCCAAATGCCGAAAAGGAGC |
| S4_5'flip_rev | GCTCCTTTTCGGGATTTGGAACAGCCTATG |
| S4_3'flip_fwd | CATACTTGTCGGCAAATGGAAAAGCCGAGC |
| S4_3'flip_rev | GCTCGGCTTTTCGATTTGCCGACAAGTATG |

The table includes the names of the oligonucleotides and their DNA sequence.

#### SUPPORTING INFORMATION TEXT

##### S1 Text. Supporting Information Methods.

###### Plasmid constructions

For pBAD33::*xre*$_{opSD}$, *P. putida* KT2440 *xre* (PP_RS12675) was amplified from pUC57::*xre-res*$^{Pp}$ using primer xre_opSD_Fw, adding an optimized SD (opSD) to the 3'-end, and primer xre_Rv. The resulting PCR product was digested with KpnI and SalI and ligated into pBAD33.

For pBAD33::*xre*$_{naSD}$-*res*$^{Pp}$, *P. putida* KT2440 *xre-res* locus (PP_RS12675- PP_RS12680) was amplified from pUC57::*xre-res*$^{Pp}$ using primer xre_naSD_Fw, including the native SD (opSD) at the 3'-end, and primer res_Rv. The resulting PCR product was digested with KpnI and SalI and ligated into pBAD33.

For pBAD33::*xre*$_{naSD}^{R67A}$-*res*$^{Pp}$, substitution of arginine-67 (R67) in Xre with alanine (A) was done using Q5 Site-Directed Mutagenesis Kit from NEB with pBAD33::*xre*$_{naSD}$-*res*$^{Pp}$ as the template and primers xre(R67A)_Fw and xre(R67A)_Rv.

*P. putida* KT2440 *xre-res* locus (PP_RS12675- PP_RS12680) was amplified from pUC57::*xre-res*$^{Pp}$ using primer xre_naSD_Fw, including the native SD (opSD) at the 3'-end, and primer res_Rv. The resulting PCR product was digested with KpnI and SalI and ligated into pBAD33.

For pGH254Kgfp, the *lacZYA* operon from pGH254K was replaced with *gfpmut2* using Gibson Assembly (NEB) according to the protocol. The *gfpmut2* gene encodes a GFP mutant containing a triple substitution; S65A, V68L, and S72A, that confers enhanced fluorescence emission and more efficient folding at 37°C(6). The vector backbone was amplified from pGH254K using primer pGH254K_gib_Fw and pGH254K_gib_Rv, while the *gfpmut2* insert was amplified from synthetic DNA ordered from Twist Bioscience. Sequencing revealed an insertion (T6613) in the ribosomal binding site (RBS), which was removed using the Q5 Site-Directed Mutagenesis Kit from NEB. Primers pGH254Kgfp_T6613_Fw and pGH254Kgfp_T6613_Rv were designed using NEBaseChanger.

For pGH254Kgfp::$P_{XR}$, The *xre-res* promoter ($P_{XR}$) was amplified from pUC57::*xre-res* using primers PXR_Fw and PXR_Rv. The resulting PCR product was digested with EcoRI and BamHI and ligated into pGH254Kgfp.

For pGH254Kgfp::$P_{XR}$ΔS1, deletion of S1 from $P_{XR}$, corresponding to basepair 1-34, was done using Q5 Site-Directed Mutagenesis Kit from NEB with pGH254Kgfp::$P_{XR}$ as the template and primers PXRΔS1_Fw and PXRΔS1_Rv.

For pGH254Kgfp::$P_{XR}$ΔS1-2, deletion of S1-2 from $P_{XR}$, corresponding to basepair 1-71, was done using Q5 Site-Directed Mutagenesis Kit from NEB with pGH254Kgfp::$P_{XR}$ as the template and primers PXRΔS1-2_Fw and PXRΔS1-2_Rv.

For pGH254Kgfp::$P_{XR}$ΔS1-3, deletion of S1-3 from $P_{XR}$, corresponding to basepair 1-101, was done using Q5 Site-Directed Mutagenesis Kit from NEB with pGH254Kgfp::$P_{XR}$ as the template and primers PXRΔS1-3_Fw and PXRΔS1-3_Rv.

For pGH254Kgfp::$P_{XR}$ΔS4, deletion of S4 from $P_{XR}$, corresponding to basepair 107-137, was done using Q5 Site-Directed Mutagenesis Kit from NEB with pGH254Kgfp::$P_{XR}$ as the template and primers PXRΔS4_Fw and PXRΔS4_Rv.

For pGH254Kgfp::$P_{XR}$ΔS3-4, deletion of S3-4 from $P_{XR}$, corresponding to basepair 71-137, was done using Q5 Site-Directed Mutagenesis Kit from NEB with pGH254Kgfp::$P_{XR}$ as the template and primers PXRΔS3-4_Fw and PXRΔS3-4_Rv.

For pGH254Kgfp::$P_{XR}$Δrepeat, deletion of sequence 3-4 from $P_{XR}$, corresponding to basepair 115-129, was done using Q5 Site-Directed Mutagenesis Kit from NEB with pGH254Kgfp::$P_{XR}$ as the template and primers PXRΔrepeat_Fw and PXRΔrepeat_Rv.

For pGH254Kgfp::$P_{XR}$-5'perfect, substitution G115T in $P_{XR}$, generating a perfect 5'-repeat in Sequence 4, was done using Q5 Site-Directed Mutagenesis Kit from NEB with pGH254Kgfp::$P_{XR}$ as the template and primers PXR-5'perf_Fw and PXR-5'perf_Rv.

For GH254Kgfp::$P_{XR}$-3'perfect, substitution A130C in $P_{XR}$, generating a perfect 3'-repeat in Sequence 4, was done using Q5 Site-Directed Mutagenesis Kit from NEB with pGH254Kgfp::$P_{XR}$ as the template and primers PXR-3'perf_Fw and PXR-3'perf_Rv.

For pGH254Kgfp::$P_{XR}$-5'flip, substitution 113-TTGTCG-118 to CGACAA in $P_{XR}$, destroying 5'-repeat in Sequence 4, was done using Q5 Site-Directed Mutagenesis Kit from NEB with pGH254Kgfp::$P_{XR}$ as the template and primers PXR-5'flip_Fw and PXR-5'flip_Rv.

For pGH254Kgfp::$P_{XR}$-3'flip, substitution 129-CGAAAA-134 to TTTTCG in $P_{XR}$, destroying 3'-repeat in Sequence 4, was done using Q5 Site-Directed Mutagenesis Kit from NEB with pGH254Kgfp::$P_{XR}$ as the template and primers PXR-3'flip_Fw and PXR-3'flip_Rv.

For pET-29b(+)::$xre_{CHis6}^{Pp}$, *P. putida* KT2440 *xre* (PP_RS12675) was designed to fuse a C-terminal $His_6$-tag directly onto the C-terminal of Xre, followed by a stop codon ($xre_{CHis6}$). The construct was custom synthesized by Twist Bioscience and cloned into pET-29b(+) using restriction sites NdeI and XhoI.

For pETDuet::$res_{NHis6}$-$xre^{Pp}$, *P. putida* KT2440 *xre* (PP_RS12675) was amplified from pUC57::*xre-res*$^{Pp}$ using primer xre_mcsII_Fw and xre_mcsII_Rv. The resulting PCR product was digested with NdeI and XhoI and ligated into MCSII of pETDuet-1 to generate pETDuet::*xre(mcsII)*. Subsequently, *P. putida* KT2440 *res* (PP_RS12680) and was amplified from pUC57::*xre-res*$^{Pp}$ using primer res_mcsI_Fw and res_mcsI_Rv. The resulting PCR product was digested with BamHI and SalI and ligated into MCSI of pETDuet::*xre(mcsII)*.

For pETDuet::$xre_{NHis6}$-$res^{Pp}$, *P. putida* KT2440 *xre* (PP_RS12675) and was amplified from pUC57::*xre-res*$^{Pp}$ using primer xre_mcsI_Fw and xre_mcsI_Rv. The resulting PCR product was digested with BamHI and SalI and ligated into MCSI of pETDuet-1 to generate pETDuet::*xre(mcsI)*. Subsequently, *P. putida* KT2440 *res* (PP_RS12680) was amplified from pUC57::*xre-res*$^{Pp}$ using primer res_mcsI_Fw and res_mcsI_Rv. The resulting PCR product was digested with NdeI and XhoI and ligated into MCSII of pETDuet::*xre(mcsI)*.

#### SUPPORTING INFORMATION REFERENCES
